## Supplementary material for "Precise optical control of gene expression in *C. elegans* using genetic code expansion and Cre recombinase": Figure Supplements

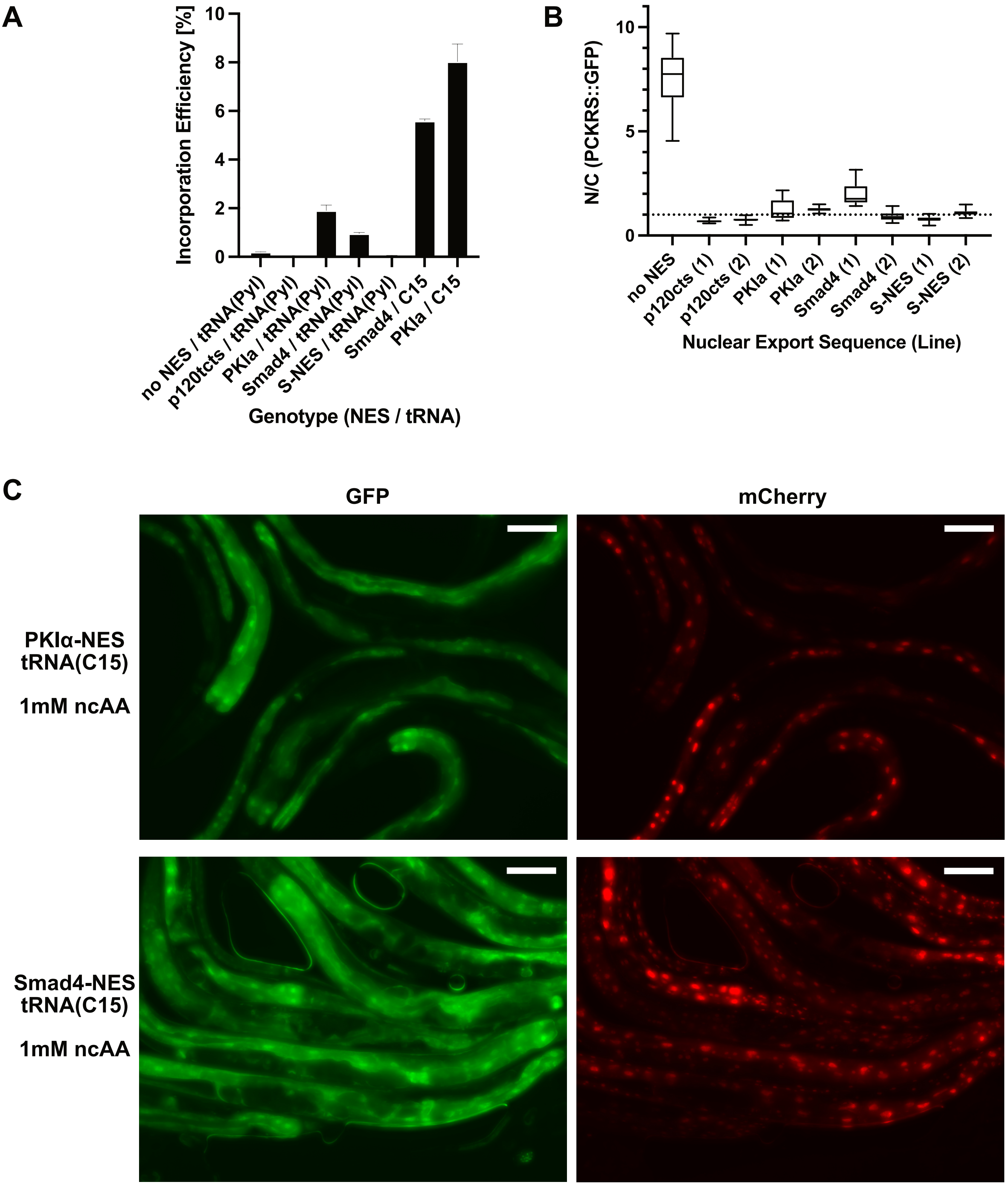


**Figure 2 - Figure Supplement 1. (A)** Quantitative Western blot of the lines blotted in Figure 2c. Each line was measured twice, the graph shows the mean and SEM. **(B)** Nuclear to cytoplasmic ratio of PCKRS::GFP fusion proteins with different nuclear export sequences for the genotypes shown in Figure 2b. Two independent lines were measured for each nuclear export sequence. N/C ratios were determined for n> 12 cells taken from at least three animals for each line. The animals shown in Figure 2b are from the first line for each NES (labeled "(1)"). Statistical significance was determined by Mann-Whitney, p<0.0001 for all NES lines compared to the wild type no NES PCKRS::GFP. **(C)** Fluorescent images of worms shown in Figure 2E. Only the panels of animals grown in the presence of 1mM ncAA are shown. GFP indicates expression of reporter construct, mCherry indicates presence of full-length reporter protein. Scale bars 30 µm.

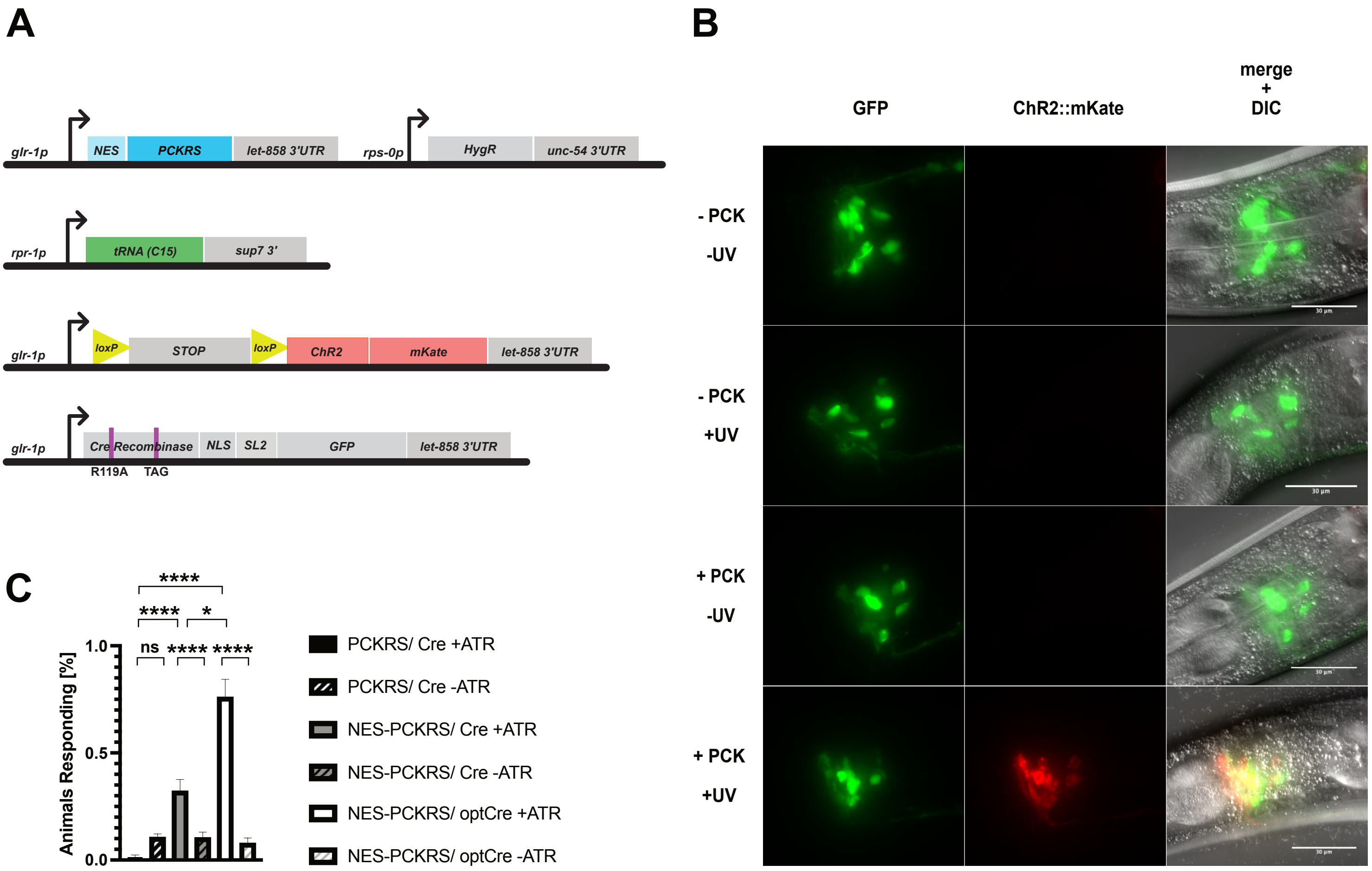


**Figure 3 - Figure Supplement 1 (A)** Constructs used to generate *glr-1p* driven photoactivatable Cre recombinase lines. **(B)** Photoactivation of P*glr-1* driven PC-Cre recombinase before optimisation, with an N-terminal NLS and intact internal NLS. Scale bars 30 µm. **(C)** Percentage of animals reversing in response to a blue light pulse for original PCKRS and *M. mazei* tRNA(Pyl)$_{CUA}$ ("PCKRS"), modified NES-PCKRS and tRNA(C15) ("NES-PCKRS"), original photocaged Cre recombinase ("PC-Cre") and optimized photo-caged Cre ("optPC-Cre"). Either in the presence ("+ATR") or absence ("-ATR") of all-trans-retinal. Significance obtained by unpaired t-test. The error is the SEM (n=4).

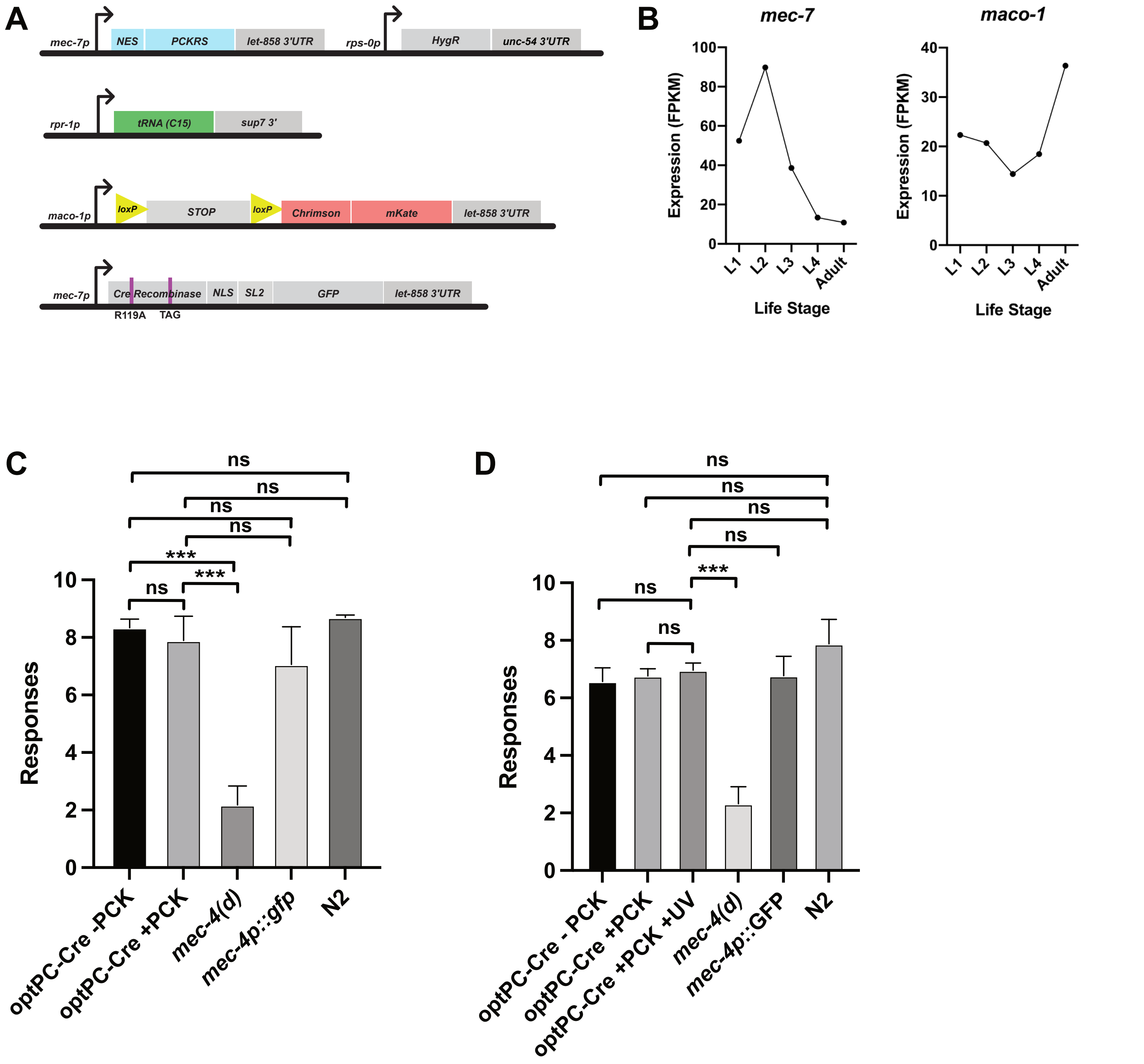


**Figure 4 - Figure Supplement 1. (A)** Constructs used to generate the *mec-7p* driven optPC-Cre recombinase lines for targeted expression of Chrimson::mKate2. **(B)** Aggregate expression estimates for *mec-7* and *maco-1* as displayed on wormbase.org. Estimates of expression calculated by averaging the FPKMs from multiple published datasets (see wormbase.org). **(C,D)** Soft Touch assays. Animals were subjected to alternating head/tail touches for a total of 10 touches. the mean of three experiments is depicted, each experiment was performed with 10 animals. **(C)** "+PCK" indicates worms grown on PCK for 48h. **(D)** "+PCK" indicates worms grown for 48h on PCK, "+UV" animals were illuminated with 365nm light at 48h. All animals were assayed 24h after the illumination illumination timepoint. Significance was determined by Mann-Whitney U test. ***p<0.001

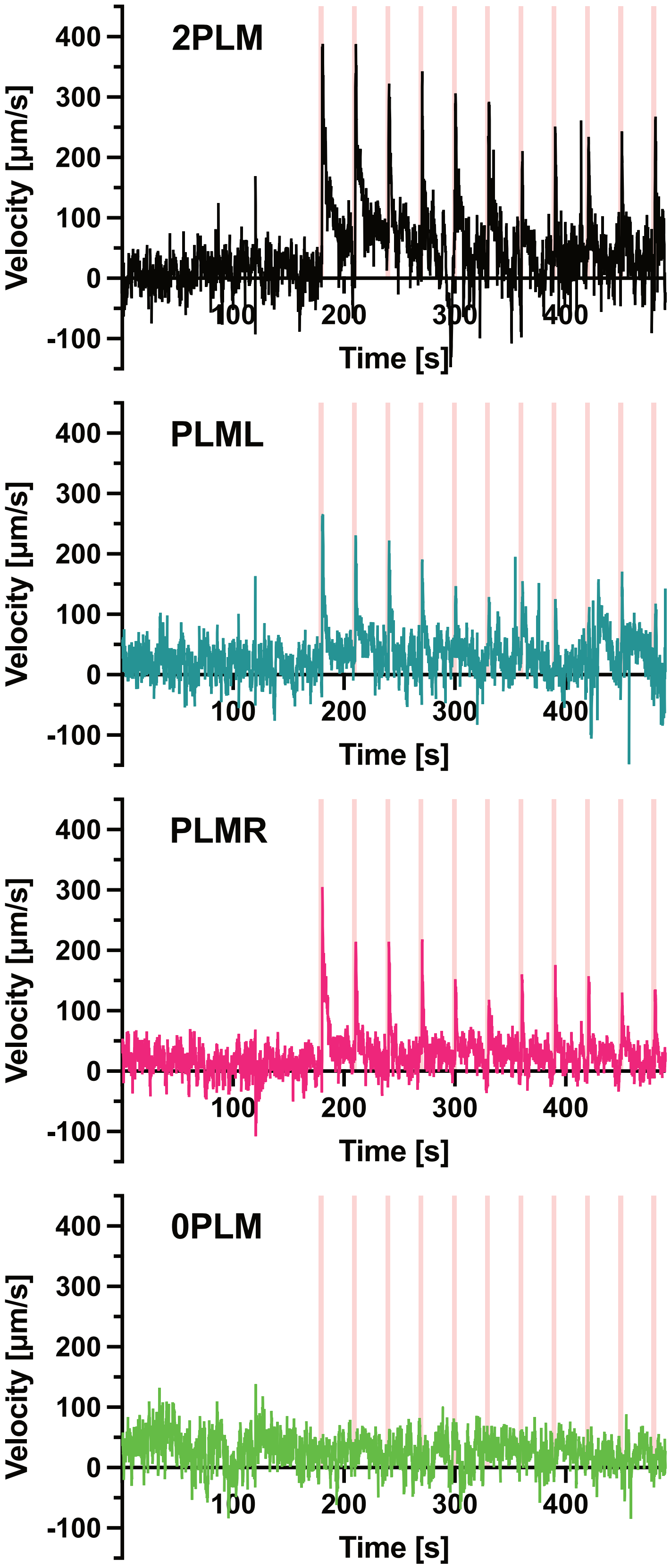


**Figure 5 - Figure Supplement 1** Speed traces used in Figure 5a. Animals expressing Chrimson::mKate2 were stimulated with 1s pulses of 617 nm light at 74 mW/cm$^2$. The pulses were delivered at 30s intervals (pink vertical lines). The assayed animals expressed Chrimson::mKate2 either in both PLM neurons (top panel), only in PLML (second panel), or only in PLMR (third panel). The bottom panel depicts mock treated animals not expressing Chrimson::mKate2. The mock treated animals were raised on PCK for 48h, then shifted to plates containing ATR for 24h before the assay was performed. The panels depict the mean velocity of 2 experiments (2PLM), 4 experiments (PLML, PLMR), and 3 experiments (0PLM). Each experiment was performed with between 4-8 animals.

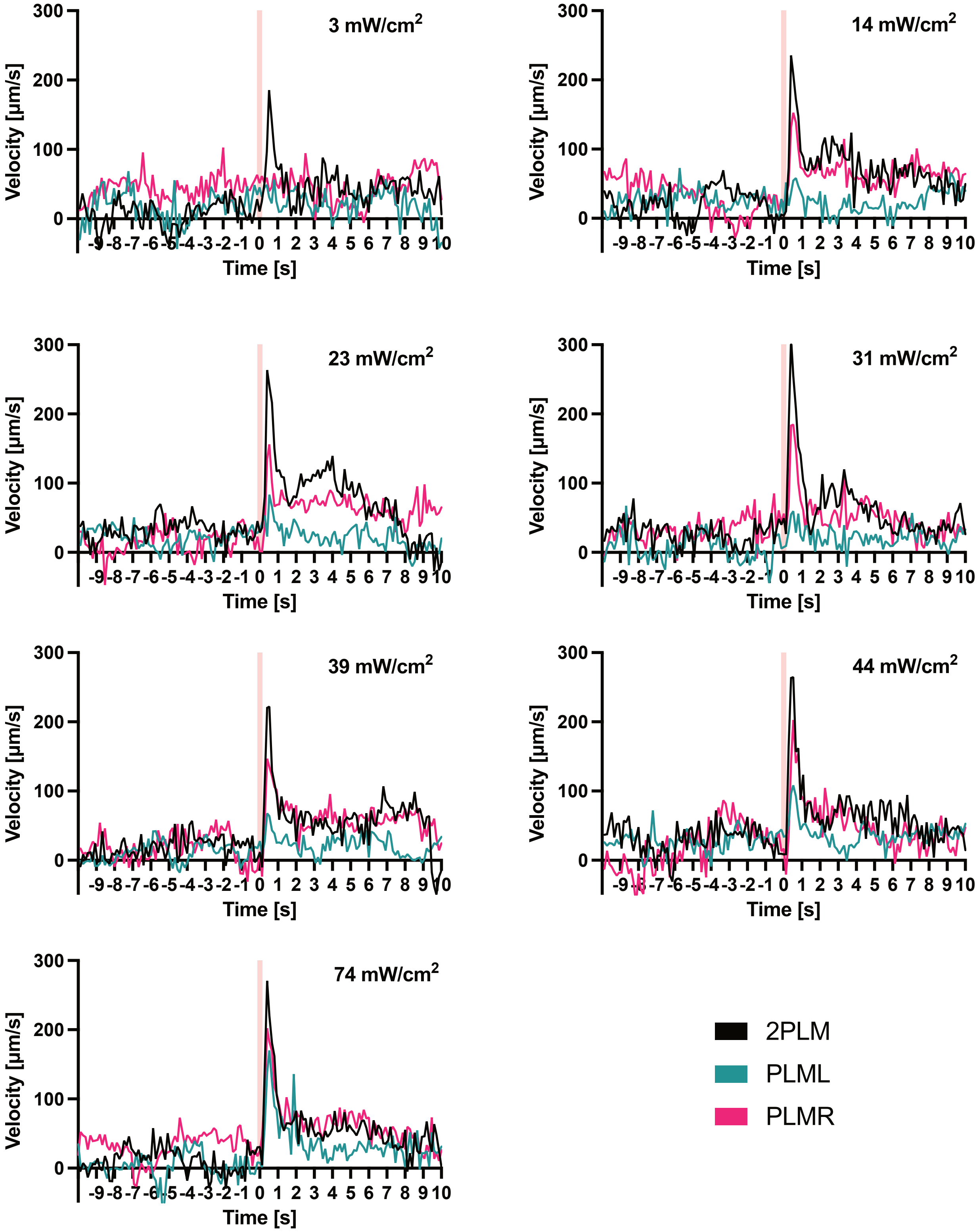


**Figure 5 - Figure Supplement 2.** Traces before normalisation of experiments shown in Figure 5C,D. Animals expressing Chrimson::mKate2 were stimulated with 0.1s pulses of 617 nm light at the indicated intensities. Pink vertical lines indicate the time point when the pulses were delivered. The panels depict the mean velocity of 3 experiments performed for each condition. Each experiment was performed with between 4 to 8 animals.

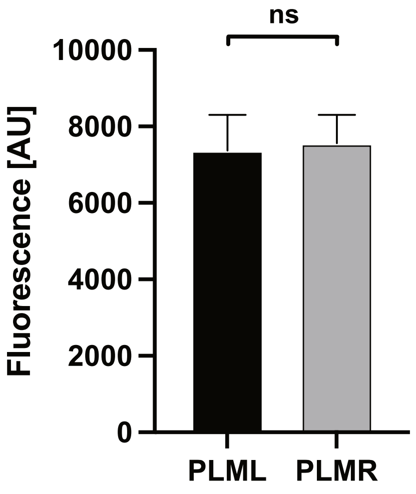


**Figure 5 - Figure Supplement 3.** Expression levels of Chrimson::mKate2. Expression levels were determined by measuring red mKate2 fluorescence in animals expressing Chrimson::mKate2 in either PLML (n=7) or PLMR (n=9). Significance was determined using the Mann-Whitney U test.
