## Supplementary Table 1 for "Precise optical control of gene expression in *C. elegans* using genetic code expansion and Cre recombinase"

### Expression Constructs

#### Destination Vectors

| Name | Description | Source |
|---|---|---|
| IR98 | *pDEST rps-0p::HygR* | [1] |
| IR226 | *glr-1p::PCKRS::let-858 3'UTR* | IR157 + SG451 + SG606 IR98 |
| IR361 | *glr-1p::loxP::B-gal 3'UTR::loxP::ChR2-mKate2::let-858 3'UTR* | IR157 + SG365 +IR355 pDEST R4-R3 II |
| LD479 | *mec-7p::Smad4 NES-PCKRS::let-858 3'UTR* | ZX204 + SE154 + SG606 IR98 |
| LD490 | *maco-1p::loxP::B-gal 3'UTR::loxP::Chrimson-mKate2::let-858 3'UTR* | SE77 + SG365 + LD287 pDEST R4-R3 II |
| SE32 | *glr-1p::Cre(R119A,K201TAG)::egl-13NLS::SL2::GFP::let-858 3'UTR* | IR157 + SE17 + IR182 pDEST R4-R3 II |
| SE163 | *sur-5p::Mm PCKRS CeOpt noStop::GFP noATG_let-858 3' UTR* | SE72 + SE156 + SG304 IR98 |
| SE164 | *sur-5p::p120cts Mm PCKRS CeOpt noStop::GFP noATG_let-858 3' UTR* | SE72 + SE157 + SG304 IR98 |
| SE165 | *sur-5p::PKIa Mm PCKRS CeOpt noStop::GFP noATG_let-858 3' UTR* | SE72 + SE158 + SG304 IR98 |
| SE166 | *sur-5p::Smad4 Mm PCKRS CeOpt noStop::GFP noATG_let-858 3' UTR* | SE72 + SE159 + SG304 IR98 |
| SE167 | *sur-5p::S NES Mm PCKRS CeOpt noStop::GFP noATG_let-858 3' UTR* | SE72 + SE160 + SG304 IR98 |
| SE168 | *sur-5p::p120cts Mm PCKRS CeOpt::let-858 3'UTR* | SE72 + SE152 + SG304 IR98 |
| SE169 | *sur-5p::PKIa Mm PCKRS CeOpt::let-858 3'UTR* | SE72 + SE153 + SG304 IR98 |
| SE170 | *sur-5p::Smad4 Mm PCKRS CeOpt::let-858 3'UTR* | SE72 + SE154 + SG304 IR98 |
| SE171 | *sur-5p::S NES Mm PCKRS CeOpt::let-858 3'UTR* | SE72 + SE155 + SG304 IR98 |
| SE174 | *glr-1p::Smad4 NES- PCKRS::let-858 3' UTR* | IR157 + SE154 + SG606 IR98 |
| SE200 | *sur-5p::Mm PCKRS CeOpt::let-858 3'UTR* | SE72 + SG451 + SG606 IR98 |
| SE284 | *mec-7p::Cre(R119A,K201TAG)::egl13NLS::SL2::GFP::let-858 3'UTR* | ZX204 + SE17 + IR182 pDEST R4-R3 II |
| SG88 | *rps-0p::GFP-mCherry-HA-NLS::unc-54 3'UTR* | [2] |
| ZS11 | *glr-1p::SV40 NLS-Cre(K201TAG)::SL2::GFP::let-858 3'UTR* | IR157 + SG617 + IR182 pDEST R4-R3 II |

#### pENTR P4-P1r Vectors

| Name | Description | Source |
|---|---|---|
| IR157 | *glr-1p* | Cloned as a P4-P1R vector using sequence from Maricq et al. [3] |
| SE72 | *sur-5p* | PCR amplified from genomic DNA with primers -<br>sur-5ps attB4F:<br>GGGGACAACTTTGTATAGAAAAGTTGCGCAGGCGGTAAACATACGTTG<br>sur-5p attB1R:<br>GGGGACTGCTTTTTTGTACAAACTTGTCTGAAAACAAAATGTAAAGTTCAAAGG |

| | | |
|---|---|---|
| SE77 | *maco-1p* | PCR amplified from genomic DNA with primers -<br>maco-1p attB4F:<br>GGGGACAACTTTGTATAGAAAAGTTGTAatttctcatgtttgttttgaaaaaaaaaccaaaag<br>maco-1p attB1R:<br>GGGGACTGCTTTTTTGTACAAACTTGTaatctgaaatacaatatatacagttatttcaatatttaacaatcaaac |
| ZX204 | *mec-7p* | PCR amplified from genomic DNA with primers -<br>mec-7p attB4F:<br>GGGGACAACTTTGTATAGAAAAGTTGTAgtttcaagatgaaacgttttgtgtgtagc<br>mec-7p attB1R:<br>GGGGACTGCTTTTTTGTACAAACTTGTcgacgtttcttcctctacacctaca |

#### pENTR 221 Vectors

| Name | Description | Source |
|---|---|---|
| SE17 | Cre(R119A,K201TAG)::egl-13NLS | Modified from SG617. The N-Terminal SV40 was removed and an egl-13 NLS was attached to the C-terminus by overlap extension PCR, A R119A mutation was inserted using primers –<br>Cre R119A F: CATGCGTgcTATCCGTAAGGAGAACGTCGACG<br>Cre R119A R: CTTACGGATAgcACGCATGACGAGGGAGACGG |
| SE150 | rpr-1p::Bt mttRNA C15::sup-7 3' | PylT in SG322 was replaced with Bt mttRNA C15<br>Sequence:<br>GGAAACCTGgTCAgGgAGAcCGAAcGGACTCTAAATCCGTTCAGCCGGGTTcGATTCCCGGGGTTTCCG |
| SE152 | p120cts NES-Mm PCKRS CeOpt | made from SG451. FLAG tag at the N-terminus was replaced by sequence encoding Human p120cts NES<br>Sequence:<br>GAGTCCCTCGAGGAGGAGCTCGACGTCCTCGTCCTCGACGACGAGGGAGGA |
| SE153 | PKIa NES-Mm PCKRS CeOpt | made from SG451. FLAG tag at the N-terminus was replaced by sequence encoding Human PKIa NES<br>Sequence:<br>CTCGCCCTCAAGCTCGCCGGACTCGACATC |
| SE154 | Smad4 NES-Mm PCKRS CeOpt | made from SG451. FLAG tag at the N-terminus was replaced by sequence encoding Human Smad4 NES<br>Sequence:<br>GCCTGCCCAGTCCCACTCCAACTCCCACCACTCGAGCGTCTCACCCTCGAC |
| SE155 | S NES-Mm PCKRS CeOpt | made from SG451. FLAG tag at the N-terminus was replaced by sequence encoding S-NES<br>Sequence:<br>GCCTGCCCAGTCCCACTCCAACTCCCACCACTCGAGCGTCTCACCCTCGAC |
| SE156 | Mm PCKRS CeOpt noStop | Stop codon was removed from SG451 using site directed mutagenesis<br>Mm PylRS noStop F: CACCAACCTCAACCCAGCTTTCTTGTACAAAGTTG<br>Mm PylRS noStop R: AAGCTGGGTTGAGGTTGGTGGAGATTCCGTTG |
| SE157 | p120cts NES-Mm PCKRS CeOpt noStop | Stop codon was removed from SG451 using site directed mutagenesis<br>Mm PylRS noStop F: CACCAACCTCAACCCAGCTTTCTTGTACAAAGTTG<br>Mm PylRS noStop R: AAGCTGGGTTGAGGTTGGTGGAGATTCCGTTG |
| SE158 | PKIa NES-Mm PCKRS CeOpt noStop | Stop codon was removed from SG451 using site directed mutagenesis<br>Mm PylRS noStop F: CACCAACCTCAACCCAGCTTTCTTGTACAAAGTTG<br>Mm PylRS noStop R: AAGCTGGGTTGAGGTTGGTGGAGATTCCGTTG |
| SE159 | Smad4 NES-Mm PCKRS CeOpt noStop | Stop codon was removed from SG451 using site directed mutagenesis<br>Mm PylRS noStop F: CACCAACCTCAACCCAGCTTTCTTGTACAAAGTTG<br>Mm PylRS noStop R: AAGCTGGGTTGAGGTTGGTGGAGATTCCGTTG |
| SE160 | S NES Mm PCKRS CeOpt noStop | Stop codon was removed from SG451 using site directed mutagenesis<br>Mm PylRS noStop F: CACCAACCTCAACCCAGCTTTCTTGTACAAAGTTG<br>Mm PylRS noStop R: AAGCTGGGTTGAGGTTGGTGGAGATTCCGTTG |

| | | |
|---|---|---|
| SG322 | rpr-1p::PylT:: sup-7 3' | rpr-1p was amplified from genomic DNA and fused to PylT by overlap extension PCR, 100bp of the sup-7 3' region was also fused to the 3' end of PylT by overlap extension PCR |
| SG365 | loxP::$\beta$-gal 3'UTR::loxP | based on Macosko et al. [4] |
| SG451 | *C. elegans* optimised M. mazei PCKRS | Synthesised after *C. elegans* optimisation[5], the gene contains mutations described in Gautier et al.[6], contains N-terminal FLAG Tag |
| SG617 | SV40 NLS-Cre(K201TAG) | Synthesised after *C. elegans* optimisation[5], inserted into pDONR221 |

#### pENTR P2R-P3 Vectors

| Name | Description | Source |
|---|---|---|
| IR182 | SL2-GFP::let-858 3'UTR | the gpd-2/3 intergenic region was amplified from genomic DNA and fused to optimised GFP. A let-858 3' UTR from SG606 was attached to the 3' end of GFP. |
| IR355 | ChR2::mKate2::let-858 3'UTR | ChR2 and mKate2 were synthesised after *C. elegans* optimisation[5], the let-858 3'UTR from SG606 was fused to the mKate2 3' end by overlap extension PCR |
| LD287 | Chrimson::mKate2::let-858 3'UTR | Chrimson was synthesised after *C. elegans* optimisation[5], mKate2::let-858 3'UTR from plasmid IR355 was fused to the 3'End of Chrimson by overlap extension PCR |
| SG304 | GFP::let-858 3'UTR | a let-858 3'UTR was fused to the 3'End of GFP by overlap extension PCR |
| SG606 | let-858 3'UTR | PCR amplified from genomic DNA<br>725 let-858 3' attB2R:<br>GGGGACCACTTTGTACAAGAAAGCTGGGTATACGGATTCGCATTTGCCAAGC<br>726 let-858 3' attB3R:<br>GGGGACAACTTTGTATAATAAAGTTGATACGGATTCGCATTTGCCAAGC |
