## Supplementary Table 2 for "Precise optical control of gene expression in *C. elegans* using genetic code expansion and Cre recombinase"

| Key Resources Table | | | | |
|---|---|---|---|---|
| **Reagent type (species) or resource** | **Designation** | **Source or reference** | **Identifiers** | **Additional information** |
| genetic reagent (*C. elegans*) | SGR30 | This paper | n/a | *greEx17[sur-5p::FLAG::PCKRS::GFP]*<br><br>Figure 2D, figure suppl 1B |
| genetic reagent (*C. elegans*) | SGR31 | This paper | n/a | *greEx18[sur-5p::p120cts::PCKRS::GFP]*<br><br>Figure 2D, figure suppl 1B |
| genetic reagent (*C. elegans*) | SGR32 | This paper | n/a | *greEx19[sur-5p::PKIα::PCKRS::GFP]*<br><br>Figure 2D, figure suppl 1B |
| genetic reagent (*C. elegans*) | SGR33 | This paper | n/a | *greEx20[sur-5p::smad-4::PCKRS::GFP]*<br><br>Figure 2D, figure suppl 1B |
| genetic reagent (*C. elegans*) | SGR34 | This paper | n/a | *greEx21[sur-5p::S-NES::PCKRS::GFP]*<br><br>Figure 2D, figure suppl 1B |
| genetic reagent (*C. elegans*) | SGR35 | This paper | n/a | *greEx22[sur-5p::PCKRS; rpr-1p::tRNA(Pyl); rps-0p::GFP(am)::mCherry::HA]; smg-2(e2008)*<br>Figure 2C,E,F,G, fig suppl 1A |
| genetic reagent (*C. elegans*) | SGR36 | This paper | n/a | *greEx23[sur-5p::p120cts::PCKRS; rpr-1p::tRNA(Pyl); rps-0p::GFP(am)::mCherry::HA];smg-2(e2008)*<br>Figure 2C, fig suppl 1A |
| genetic reagent (*C. elegans*) | SGR37 | This paper | n/a | *greEx24[sur-5p::PKIα::PCKRS; rpr-1p::tRNA(Pyl); rps-0p::GFP(am)::mCherry::HA]; smg-2(e2008)*<br>Figure 2C, fig suppl 1A |
| genetic reagent (*C. elegans*) | SGR38 | This paper | n/a | *greEx25[sur-5p::smad-4::PCKRS; rpr-1p::tRNA(Pyl); rps-0p::GFP(am)::mCherry::HA]; smg-2(e2008)* |

| | | | | |
|---|---|---|---|---|
| | | | | Figure 2C, fig suppl 1A |
| genetic reagent (*C. elegans*) | SGR39 | This paper | n/a | *greEx26[sur-5p::SNES::PCKRS; rpr-1p::tRNA(Pyl); rps-0p::GFP(am)::mCherry::HA]; smg-2(e2008)* Figure 2C, fig suppl 1A |
| genetic reagent (*C. elegans*) | SGR40 | This paper | n/a | *greEx27[sur-5p::PCKRS; rpr-1p::tRNA(Pyl); rps-0p::GFP(am)::mCherry::HA]; smg-2(e2008)* Figure 2F, G |
| genetic reagent (*C. elegans*) | SGR45 | This paper | n/a | *greEx32[sur-5p::PKIα::PCKRS; rpr-1p::tRNA(C15); rps-0p::GFP(am)::mCherry::HA]; smg-2(e2008)* Figure 2C,E,F,G, fig. suppl. 1A,C |
| genetic reagent (*C. elegans*) | SGR46 | This paper | n/a | *greEx33[sur-5p::smad-4::PCKRS; rpr-1p::tRNA(C15); rps-0p::GFP(am)::mCherry::HA]; smg-2(e2008)* Figure 2C,E,F,G, fig. suppl. 1A,C |
| genetic reagent (*C. elegans*) | SGR47 | This paper | n/a | *greEx34[sur-5p:: PKIα::PCKRS; rpr-1p::tRNA(C15); rps-0p::GFP(am)::mCherry::HA];smg-2(e2008)* Figure 2F,G |
| genetic reagent (*C. elegans*) | SGR48 | This paper | n/a | *greEx35[sur-5p::smad-4::PCKRS; rpr-1p::tRNA(C15); rps-0p::GFP(am)::mCherry::HA] ;smg-2(e2008)* Figure 2F,G |
| genetic reagent (*C. elegans*) | SGR49 | This paper | n/a | *greEx36[glr-1p::PCKRS rpr-1p::tRNA(Pyl); glr-1p::PC-Cre; glr-1p::B-gal terminator + loxP::ChR2::mKate2]* Figure 3D,G, fig suppl 1C |
| genetic reagent (*C. elegans*) | SGR50 | This paper | n/a | *greEx37[glr-1p::PCKRS; rpr-1p::tRNA(Pyl); glr-1p::PC-Cre; glr-1p::B-gal terminator + loxP::ChR2::mKate2]* Figure 3D,G, fig suppl 1C |

| | | | | |
|---|---|---|---|---|
| genetic reagent (*C. elegans*) | SGR51 | This paper | n/a | *greEx38[glr-1p::smad-4::PCKRS; rpr-1p::tRNA(C15); glr-1p::PC-Cre; glr-1p::B-gal terminator + loxP::ChR2::mKate2]*<br>Figure 3D,G, fig suppl 1B,C |
| genetic reagent (*C. elegans*) | SGR52 | This paper | n/a | *greEx39[glr-1p::smad-4::PCKRS; rpr-1p::tRNA(C15); glr-1p::PC-Cre; glr-1p::B-gal terminator + loxP::ChR2::mKate2]*<br>Figure 3D,G, fig suppl 1C |
| genetic reagent (*C. elegans*) | SGR53 | This paper | n/a | *greEx40[glr-1p::smad-4::PCKRS; rpr-1p::tRNA(C15); glr-1p::optPC-Cre; glr-1p::B-gal terminator + loxP::ChR2::mKate2]*<br>Figure 3D,F,G, fig suppl 1C, suppl videos 1,2 |
| genetic reagent (*C. elegans*) | SGR54 | This paper | n/a | *greEx41[glr-1p::smad-4::PCKRS; rpr-1p::tRNA(C15); glr-1p::optPC-Cre; glr-1p::B-gal terminator +* loxP::ChR2::mKate2]<br>Figure 3D,G, fig suppl 1C |
| genetic reagent (*C. elegans*) | SGR55 | This paper | n/a | *greEx42[mec-7p::smad-4::PCKRS; rpr-1p::tRNA(C15); mec-7p::optPC-Cre; Pmaco-1::B-gal terminator + loxP::ChR2::mKate2]* |
| genetic reagent (*C. elegans*) | SGR56 | This paper | n/a | *greIs1[mec-7p::smad-4::PCKRS; rpr-1p::tRNA(C15); mec-7p::optPC-Cre; Pmaco-1::terminator + loxP::ChR2::mKate2]*<br>Fig.4C,D,E<br>Was generated by gamma integration from SGR55, backcrossed 2x |
| genetic reagent (*C. elegans*) | SGR96 | This paper | n/a | *greIs1[mec-7p::smad-4::PCKRS; rpr-1p::tRNA(C15); mec-7p::optPC-Cre; Pmaco-1::terminator + loxP::ChR2::mKate2]*<br>Figure 4 - supplement 1C,D; Figure 5, fig.suppl 1,2,3 |
| genetic reagent (*C. elegans*) | SGR97 | This paper | n/a | *greEx17[sur-5p::FLAG::PCKRS::GFP]*<br><br>Figure 2 - Supplement 1B |

| | | | | |
|---|---|---|---|---|
| genetic reagent (*C. elegans*) | SGR98 | This paper | n/a | *greEx18[sur-5p::p120cts::PCKRS::GFP]*<br><br>Figure 2 - Supplement 1B |
| genetic reagent (*C. elegans*) | SGR99 | This paper | n/a | *greEx19[sur-5p::PKIα::PCKRS::GFP]*<br><br>Figure 2 - Supplement 1B |
| genetic reagent (*C. elegans*) | SGR100 | This paper | n/a | *greEx20[sur-5p::smad-4::PCKRS::GFP]*<br><br>Figure 2 - Supplement 1B |
| genetic reagent (*C. elegans*) | SGR101 | This paper | n/a | *greEx21[sur-5p::S-NES::PCKRS::GFP]*<br><br>*Figure 2 - Supplement 1B* |
| strain, strain background (*C. elegans*) | N2 | CGC | WBStrain00000001 | wild type<br><br>Figure 4 - Supplement 1C |
| genetic reagent (*C. elegans*) | CZ10175 | CGC | WBStrain00005421 | *zdIs5[mec-4p::GFP + lin-15(+)]*<br><br>Figure 4 - Supplement 1C *("mec-4p::gfp")* |
| genetic reagent (*C. elegans*) | | Chalfie & Au 1989 (doi: 10.1126/science.2646709) | WBVar00266589 | *mec-4(u231)*<br><br>Figure 4 - Supplement 1C *("mec-4(d)")* |
| antibody | mouse anti-GFP, monoclonal (clones 7.1 &13.1) | Roche | Cat# 11814460001 | Figure 2C,F,G, fig suppl 1A (1:4000) |
| antibody | rat anti-HA, monoclonal (clone 3F10) | Roche | Cat# 11867423001 | Figure 2C,G (1:2000) |
| antibody | Horse anti-mouse IgG HRP | Cell Signaling Technology | Cat# 7076S | Figure 2C,F,G, fig suppl 1A (1:5000) |
| antibody | Goat anti-Rat IgG (H+L) HRP | Thermo Fisher Scientific | Cat# 31470 | Figure 2C,G (1:5000) |
| chemical compound | Photocaged lysine (6-nitropiperonyl | ChiroBlock GmbH;<br><br>Gaultier et at, 2010 | n/a | custom synthesised by ChiroBlock GmbH, Germany.<br><br>Synthesis described in |

| | | | | |
|---|---|---|---|---|
| | -L-Lysine) | | | Gautier et al. (doi: 10.1021/ja910688s) |
